# Role of Diversity-Generating Retroelements for Regulatory Pathway Tuning in Cyanobacteria

**DOI:** 10.1101/2020.05.26.117283

**Authors:** Alec Vallota-Eastman, Eleanor C. Arrington, Siobhan Meeken, Simon Roux, Krishna Dasari, Sydney Rosen, Jeff F. Miller, David L. Valentine, Blair G. Paul

## Abstract

**Background:** Cyanobacteria maintain extensive repertoires of regulatory genes that are vital for adaptation to environmental stress. Some cyanobacterial genomes have been noted to encode diversity-generating retroelements (DGRs), which promote protein hypervariation through localized retrohoming and codon rewriting in target genes. Past research has shown DGRs to mainly diversify proteins involved in cell-cell attachment or viral-host attachment within viral, bacterial, and archaeal lineages. However, these elements may be critical in driving variation for proteins involved in other core cellular processes.

**Results:** Members of 31 cyanobacterial genera encode at least one DGR, and together, their retroelements form a monophyletic clade of closely-related reverse transcriptases. This class of retroelements diversifies target proteins with unique domain architectures: modular ligand-binding domains often paired with a second domain that is linked to signal response or regulation. Comparative analysis indicates recent intragenomic duplication of DGR targets as paralogs, but also apparent intergenomic exchange of DGR components. The prevalence of DGRs and the paralogs of their targets is disproportionately high among colonial and filamentous strains of cyanobacteria.

**Conclusion:** We find that colonial and filamentous cyanobacteria have recruited DGRs to optimize a ligand-binding module for apparent function in signal response or regulation. These represent a unique class of hypervariable proteins, which might offer cyanobacteria a form of plasticity to adapt to environmental stress. This analysis supports the hypothesis that DGR-driven mutation modulates signaling and regulatory networks in cyanobacteria, suggestive of a new framework for the utility of localized genetic hypervariation.

## Background

Cyanobacteria are a remarkably diverse lineage, in terms of metabolisms, morphologies, and habitat distribution. Perhaps most notably, this phylum contains the only prokaryotic organisms known to have evolved the capability for oxygenic photosynthesis; this trait was later acquired by eukaryotes through endosymbiosis with cyanobacteria, resulting in the formation of chloroplasts [1,2], and driving the modern biosphere. Cyanobacteria have evolved an array of morphologies, including complex multicellular forms [3–6]. Representatives are typically classified into five subsections [7,8]. Species of subsections I and II consist of single coccoid cells. Subsections III-V represent multicellular species that form filaments of varying complexity. Members of subsection III form reversibly-differentiable filaments of vegetative cells. Among subsections IV and V, cells can carry out terminal cellular differentiation in response to environmental stimuli, forming spore-like cells that are resistant to desiccation (akinetes), micro-oxic cells specialized for N_2_ fixation (heterocysts), and motile filaments (hormogonia) [9]. This morphological and metabolic complexity has allowed cyanobacteria to inhabit diverse environments.

Certain members of the cyanobacterial phylum possess an extensive capacity to adapt to various environmental pressures through tightly-controlled regulation of complex cellular programs for signal response. This is exemplified by abilities for metabolic switching (i.e. CO_2_/N_2_ fixation), maintaining photoreceptors of varying intensity for binary programs of circadian rhythm, and forming specialized cells which can sometimes be terminally differentiated and lead to multicellularity [9]. To regulate these complex programs, cyanobacteria have an extensive repertoire of genes governing signal transduction including proteases, kinases, and nucleases. Notably, paralogs of these regulatory proteins are more abundant among the more complex species of cyanobacteria (i.e. those belonging to subsections III-V) [10,11]. However, the mechanisms to diversify and adapt specific functionality in these duplicated genes remain largely unexplored. One mechanism may be the application of diversity-generating retroelements, known to accelerate the evolution of the proteins they target.

Diversity-generating retroelements (DGRs) have been noted within the genomes of several genera of cyanobacteria [12–14]. In other bacterial and viral systems, DGRs drive site-specific hypermutation of a subset of codons in target genes. These retroelements utilize a uniquely targeted form of retrotransposition. To this end, DGRs insert variants into a flexible coding scaffold, while avoiding non-specific variation in conserved portions of a gene. The essential features of a DGR are most often found within a single genomic locus spanning ~ 5 – 10 kbp (Fig. 1a), though the synteny and organization of DGR components can vary [13]. Diversification is mechanistically carried out by a reverse transcriptase (RT), which acts upon an RNA transcript, encoded by a template repeat (TR) region in the locus. This region is nearly identical to a variable region (VR) that typically resides in a nearby gene, which encodes a DGR-variable protein (VP). The TR-RNA intermediate is reverse transcribed into cDNA wherein A → N mutation is highly favored by the error-prone RT. This cDNA then replaces VR, whose sequence commonly corresponds to flexible residues in ligand binding structural domains belonging to the C-type lectin or Immunoglobulin (Ig)-like protein families.

**Figure 1.**
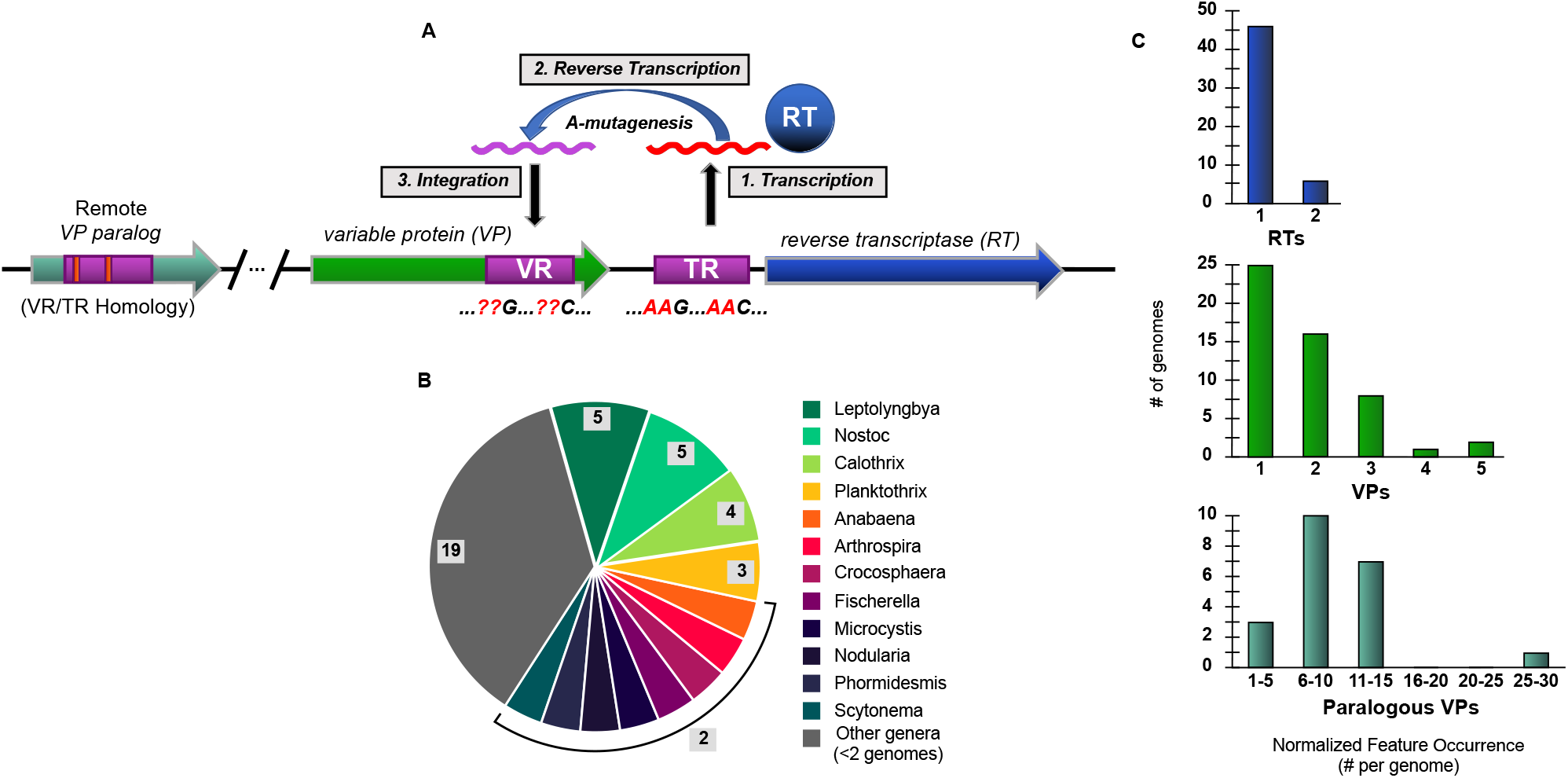
Schematic overview of a DGR and prevalence in cyanobacterial genomes. **a** Three primary steps in the process of mutagenic homing are shown: **1)** conserved template region (TR) in the DGR cassette is transcribed into intermediate, non-coding RNA, which is the substrate for DGR reverse transcriptase (DGR-RT). **2)** Template-primed reverse transcription of TR-RNA is highly error-prone at adenines, which thus incorporates random nucleotides at specific positions in the resulting cDNA. **3)** The new cDNA molecule is integrated into the variable region (VR) at a fixed locus, resulting in the replacement of a portion of the target gene (~100 – 200bp). Genomic surveys suggest that VRs occur almost exclusively near the 3’ terminus of a target gene. Additional “remote” VP genes (i.e. paralogs) may be found in non-DGR loci throughout the genome, which have detectable TR vs VR homology. **b** Summary of 52 cyanobacteria genomes known to have DGR components (in Fig. 1a) spanning 31 genera. Genera with ≥ 2 DGR-containing genomes annotated. **c** DGR feature occurrence normalized to genome number.

The first DGR variable protein was characterized from the bacteriophage, BPP-1. In these phage, DGRs diversify tail fiber tip proteins that recognize and bind to *Bordetella* host receptors [12,15]. Other cellular DGRs have been characterized in bacterial pathogens, including *Legionella pneumophila* [16] and *Treponema denticola* [17], where DGR target genes encode for cellular surface proteins, presumably involved in cell-cell attachment. The conserved function of cell-cell or viral-cell attachment in these target genes lends to a perspective of DGRs for broad use in host recognition for symbiosis or infection. Moreover, several genera of cyanobacteria were identified in recent genomic and metagenomic surveys of DGRs [13,18]. The essential components of DGRs can be found across most lineages of prokaryotic life [13,18–22], suggesting broad utility of this form of localized mutation.

Whereas previously characterized DGR target proteins appear to share a functional role in extracellular attachment to ligands displayed on foreign cells, these retroelements could potentially diversify other cellular proteins with entirely distinct functions. The intermediate RNA, which preserves a template for DGR mutagenesis, has been shown to be highly expressed in lab isolates of *Trichodesmium erythraeum* IMS101 [14] and in *Nodularia spumigena* CCY9414 under light and oxidative stress [23,24]. Here, a systematic analysis of DGRs and their variable proteins in cyanobacterial genomes leads to a new perspective on the utility of diversification and optimization of modular protein domains in paralogs that appear linked to signaling and transcriptional control.

## Results and discussion

### A Conserved Subclass of Retroelements in Cyanobacteria

Our analysis identified 58 DGRs that include 90 target genes (i.e. encoding VPs) in 52 genomes of cyanobacteria spanning 31 different genera. These include filamentous, colonial, and symbiotic organisms (Fig. 1b and Additional file 1: Table S1). Sequence clustering of the 58 DGRs (RT sequences at 95% identity) suggests they represented 49 distinct DGRs, although the full set of 58 were examined further. All DGRs were identified by presence of diagnostic and essential components: an RT gene; one or more VP genes with VR regions; and a TR region. Among the 52 genomes analyzed, four contain duplicate DGR cassettes, based on clustering, while one contains two unique DGR-RTs. Moreover, several individual DGRs have multiple target genes, and some VP genes have VRs with homology to other genes dispersed throughout the genome (paralogs) (Fig. 1c).

To evaluate the diversity of cyanobacteria-encoded DGRs, we first compared these representatives to a recently developed, global metagenomic DGR dataset [18]. Cyanobacterial DGR-RTs were clustered (i.e. at ≥ 50% AAI) with sequences in the global metagenomic dataset, then linked to a corresponding DGR clade and target protein cluster. All DGRs from our dataset were closely related to DGR Clade-5. The global dataset RTs in DGR Clade-5 are affiliated with target proteins in protein cluster 1 (i.e. PC_00001), which primarily contains cellular proteins that appear to be membrane-bound [18]. Given that the cyanobacterial DGRs appear to cluster tightly together, we next sought to analyze phylogenetic relationships within this set.

Phylogenetic analysis of cyanobacterial DGR-RTs revealed a monophyletic clade, unique from all other bacterial DGR-RTs (Fig. 2). The cyanobacterial DGR-RT clade comprises sequences that span nearly all major cyanobacterial genera within morphological subclasses I, III, IV, and V (Fig. 3a). None of the DGR-containing genomes correspond to genera within subclass II. Strikingly, cyanobacterial reverse transcriptases within the monophyletic cyanobacterial DGR clade share an average global sequence identity of 67% (minimum 55%; amino acid sequence). Whereas members of this DGR-RT subgroup do not appear to be shared with other bacteria or archaea, their phylogenetic relationships suggest a complex evolutionary history punctuated by horizontal exchange within the cyanobacterial phylum (Fig. 3b). Although none of the cyanobacterial DGRs could be definitively assigned to prophage elements, they were identified on plasmids of *Anabaena* sp. 90 (CP003287) and *Fischerella* sp. NIES-4106 (AP018301), which may indicate a vehicle for retroelement transfer between closely related populations. Among members of this RT clade, each corresponding DGR-VP contains a ligand-binding C-type lectin-like domain (CLec) with additional functional domains described in detail below.

**Figure 2.**
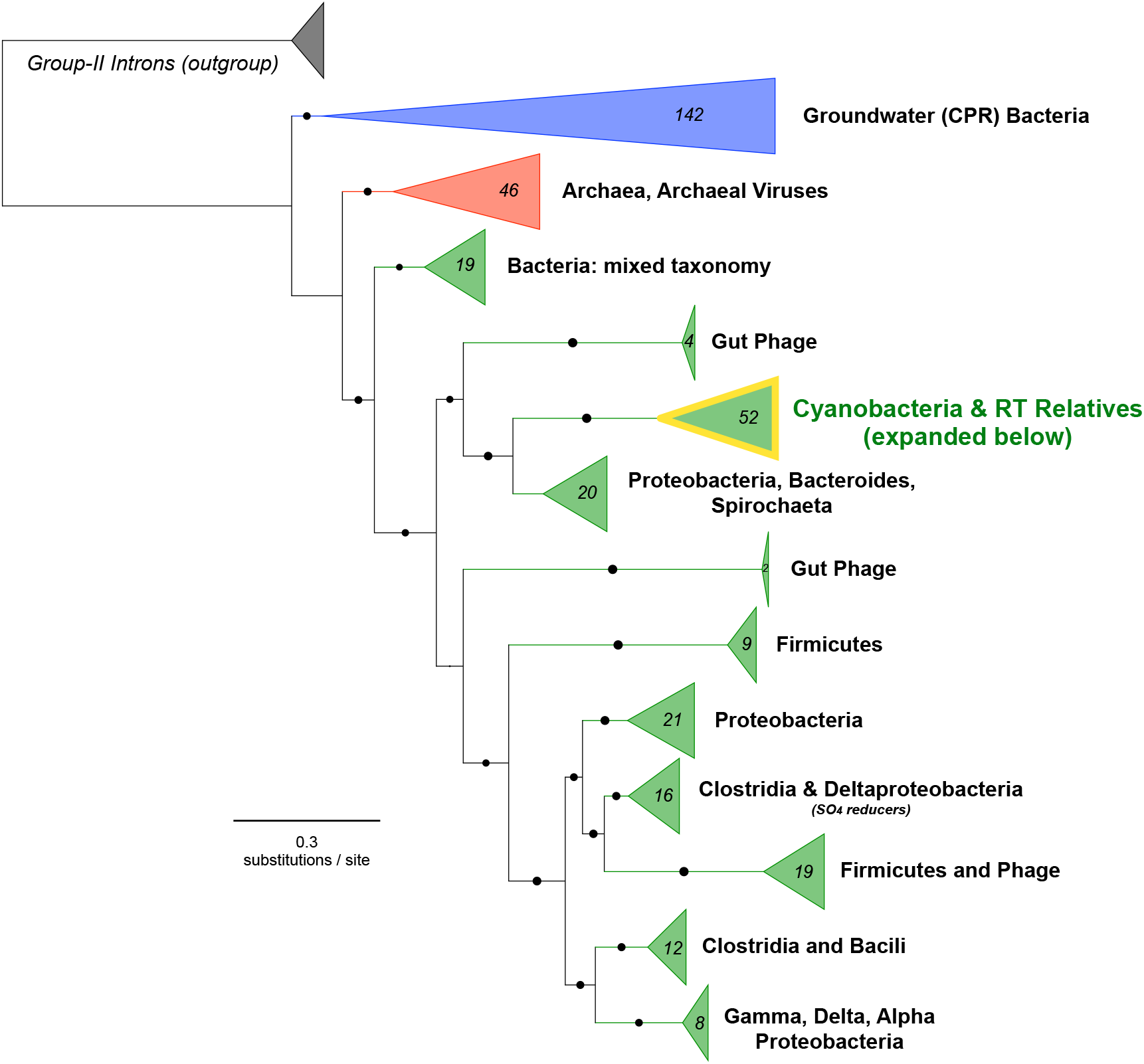
Broad RT phylogeny compared amongst all known DGR-containing lineages. RT phylogeny compared amongst all known DGR-containing lineages with Group-II Introns as the outgroup (highlighted cyanobacterial clade expanded in Figure 3).

**Figure 3.**
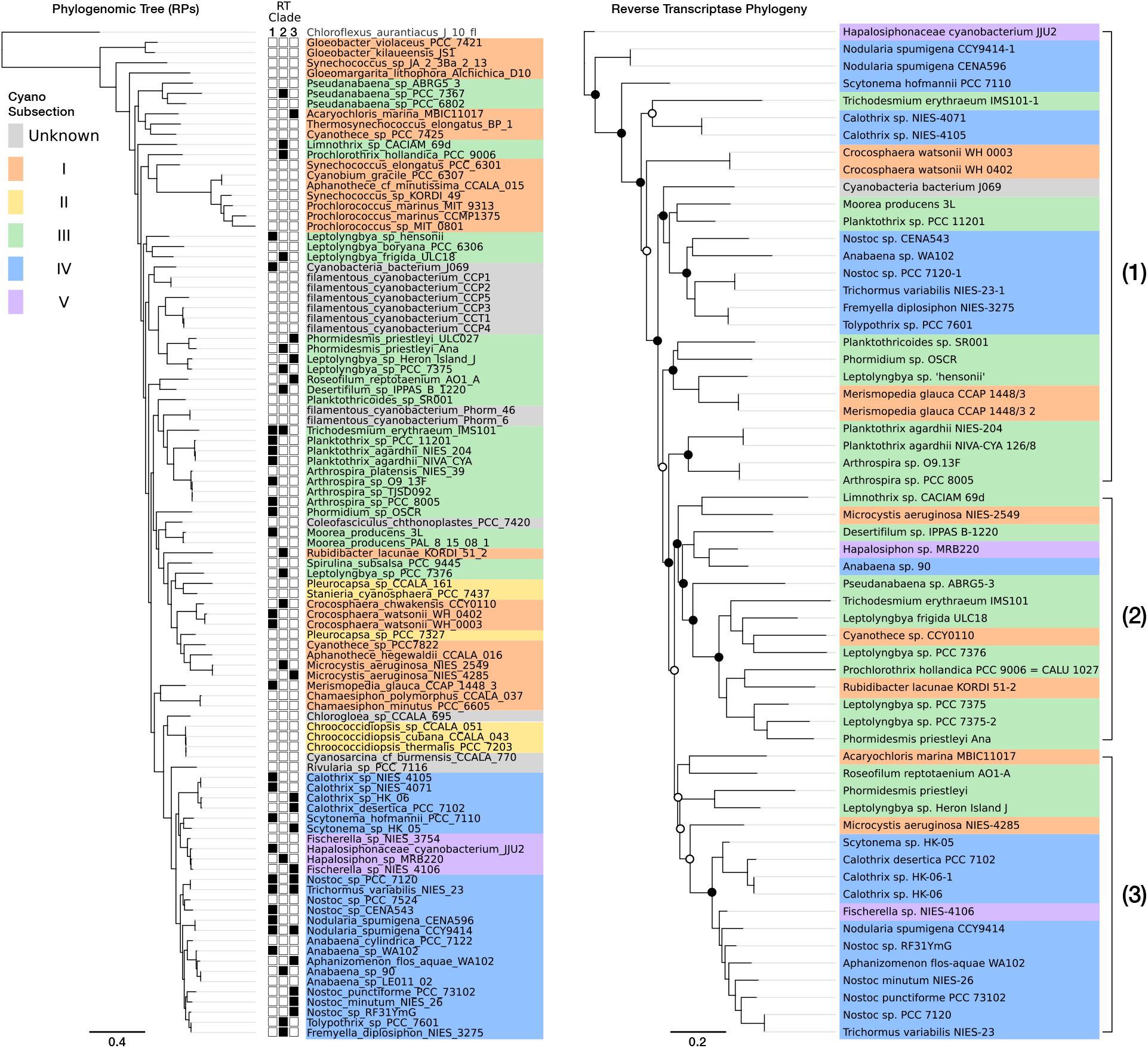
Phylogenetic reconstruction for DGR-containing cyanobacteria and DGR-RT phylogeny. **a** Phylogeny for concatenated ribosomal protein alignments, including all DGR-containing species. Filled boxes (left) indicate DGR-RT containing species and the corresponding RT clade. **b** DGR-RT phylogeny with cyanobacterial physiological subsections highlighted in color. Circles indicate branch support values (hollow >50%; filled >70%).

### Intragenomic Dispersal of Hypervariable Binding Domains

DGR variable proteins often contain multiple distinct structural domains [13,21,25]. To investigate the specific functions of cyanobacterial DGR-targeted proteins (i.e. containing the VR scaffold), we first separately analyzed the ligand-binding CLec domains in all DGR-VPs. This approach identified a conserved module (i.e. a putative C-terminal domain) found in each of the 52 cyanobacterial VP representatives (Additional file 1: Table S1). The entire set of VR-containing modules share sequence homology with 50.5% average identity and, moreover, all of these protein sequences were clustered together with >30% pairwise amino acid identity. Structural prediction of the representative C-terminal domain sequence (i.e. obtained from clustering) determined that each module most closely resembles the C-type Lectin fold, which is represented by the CLec-like superfamily (InterPro: IPR016187). A search for similar proteins in the Uniprot database identified sequences from an array of other genomes among which 92% belong to cyanobacterial phyla (Additional file 2: Table S2). The similarity between CLec domains found in diverse DGRs may underlie a conserved utility for diversifying this module across different cyanobacterial taxa. The CLec-like superfamily has been linked to a variety of molecular processes in cells and viruses spanning the tree of life, with a common functional role in ligand binding generally predicted for this fold [26–28]. Thus, the modular and dispersed nature of a highly conserved CLec subclass may further point to multifaceted functional significance in cyanobacteria.

We next sought to address whether hypervariable CLec modules might arise from gene duplication and intragenomic dispersal, resulting in recognizable sets of paralogs in cyanobacterial genomes. This search was limited to 21 high-quality genomes of the 52-genome total, such that draft genomes composed of >50 scaffolds were removed from the analysis. This approach uncovered 21 genomes that have multiple genes encoding CLec domain-containing proteins, with varying degrees of VR/TR homology (Fig. 4 and Additional file 3: Table S3). These paralogs occur both within DGR loci and dispersed throughout the genome and most often consist of either a single CLec domain or the C-terminal CLec grafted to an N-terminal putative serine kinase domain. Taken together, the multi-genome set of 219 cyanobacterial orthologs across 21 genomes–share average pairwise identity of 50.5% within their CLec domains. The complete set of 219 orthologs comprises 121 genes that appear to be DGR-diversified based on VR/TR homology, including 45 VP genes encoded within a DGR. The additional 76 remote targets were associated with their respective genome’s DGR(s) using a threshold of TR identity greater than 50%; these matches were exclusively found near the 3’ terminus of CLec-encoding genes. The proximity to 3’-termini suggests that conserved, cis-acting features - such as DNA cruciforms or initiation of mutagenic homing sites required for cDNA integration - may play a role in activating remote targets.

**Figure 4.**
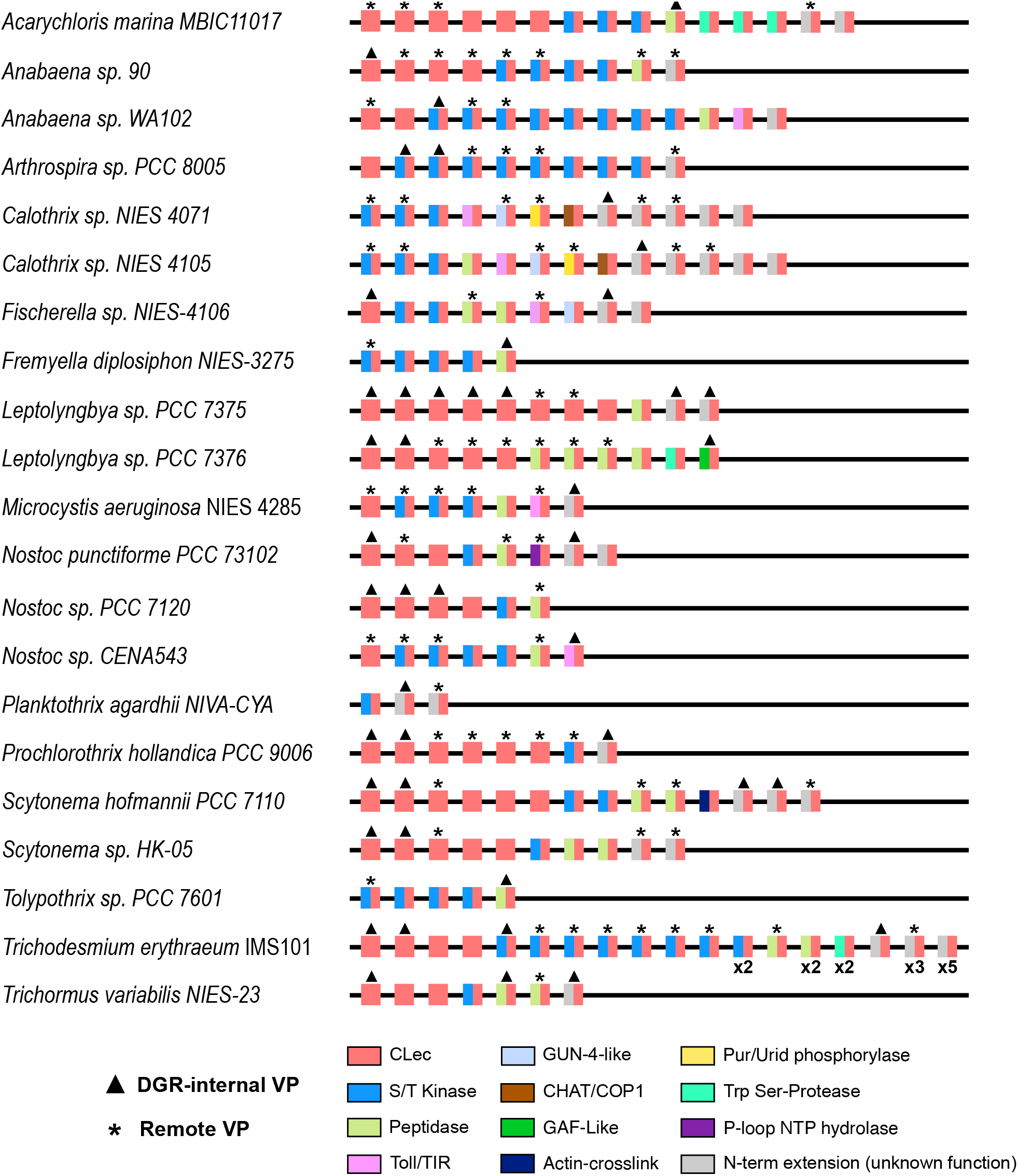
Distribution of CLec variable protein paralogs in cyanobacterial genomes. DGR variable proteins and remote variable proteins are indicated by triangles and asterisks over corresponding representatives. The CLec paralogs of VPs and RVPs are also shown for each genome. The CLec domain found in all paralogs is indicated by either a red square (CLec only) or two grafted domains shown as adjacent rectangles (C-terminal CLec in red; N-terminal domains in various colors). For clarity, paralogs of the same domain architecture in *Trichodesmium erythraeum* IMS101 are indicated below the representative protein (e.g. x2). Note: only a representative subset of DGR-containing genomes is shown (15 of 52 genomes).

The genome of *Nostoc* sp. PCC 7120 (formerly *Anabaena*), contains two DGRs and several dispersed VP paralogs (Fig. 4), providing the opportunity to examine the evolutionary history of these genes in an extensively-studied model organism. Within this genome, we identified three highly similar VP homologs (≥ 60% amino acid identity) in dispersed loci, wherein these genes may have proliferated by duplication and transposition from a common ancestral gene. Notably, one of these paralogs (*all3226*) contains remnant TR-VR homology, despite an absence of proximal RT genes or pseudogenes. Taken together, this suggests a capacity for intragenomic dispersal of DGR-targeted variable proteins, and perhaps removal of diversification components once an optimal variant is selected. In addition to its tractability, the common constellation of DGR VPs that occurs in PCC 7120, as observed in other cyanobacteria, make this species an ideal representative for further analysis of the physiological, ecological and evolutionary ramifications of DGR VP functionality and modularity in cyanobacteria.

To assess whether transposable elements were found in proximity to DGRs, we analyzed neighborhoods surrounding each hypervariable protein, including remote VPs with respect to a DGR-RT (i.e. > 5 kbp upstream/downstream). This search uncovered transposase genes belonging to various families in DGR-proximal loci which may be responsible for VP dispersal throughout the genome (Additional file 4: Table S4 and Additional file 5: Table S5). Within the subset of 21 high-quality genomes, *Trichodesmium erythraeum* IMS101 has the greatest number of proximal transposase genes, spanning six different insertion sequence (IS) families. The most widely-distributed transposases were those belonging to the IS200/IS605 family, found nearby 9 VPs from 6 distinct species. Transposases belonging to this family employ a single-stranded DNA intermediate for a “peel-and-paste” mechanism of transposition [29,30]. The genome of *Anabaena* sp. Strain 90 contains remnants of a putative degraded DGR cassette – containing only the RT with no other detectable features – and notably, the RT gene is flanked by proximal transposase genes. This provides a potential mechanism for select components of the DGR to be mobilized within the genome. DGR recruitment to one gene from another would allow favorably diversified genes to become conserved while targeting hypervariation elsewhere in the genome. Selective pressures can then influence the recruitment of DGRs to genes wherein hypervariation for ligand-binding residues offers selective advantages. Through this mechanism of transposition, cyanobacterial DGRs may provide a newly-diversified, modular, ligand binding domain to signaling genes.

### Function of Multidomain Variable Proteins

In part, functional diversity of DGR variable proteins is found in their multidomain complexity. We examined cyanobacterial VPs and their paralogs, which consist of N-terminal domains that are grafted to the C-terminal CLec domain (Fig. 4, Fig. 5). Toward assessing cellular localization, transmembrane and/or signal peptide regions were predicted for 4 DGR-associated VPs and 9 remote VPs, spanning 11 of the 21 high-quality genome set (Additional file 3: Table S3). Most cyanobacterial DGR VPs are predicted to be cytosolic, however evidence exists for TM localization and secretion as well.

**Figure 5.**
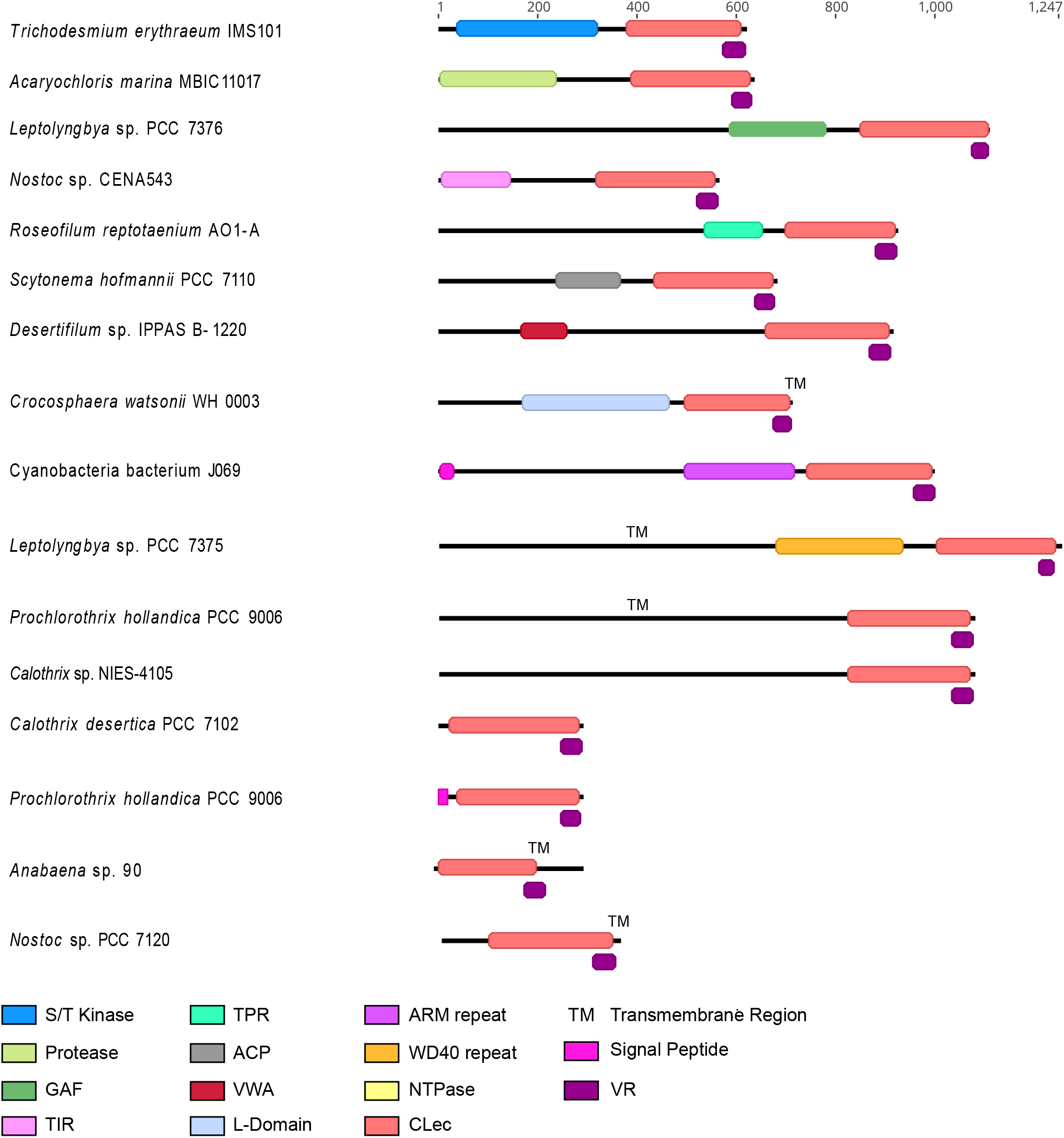
Representative variable protein (VP) domain architectures. Protein domains are colored according to pHMMR domain assignment. Variable regions are shown in deep purple. Additional features, including predicted signal peptides and transmembrane helices, are also indicated. An example species is shown to left of each VP architecture.

The most common N-terminal domains of cyanobacterial VPs and their paralogs have similarity to the protein kinase superfamily. These proteins are further predicted to be serine/threonine kinases (STKs) based on the following factors: 1) identification of Hanks and Hunter-type Motifs I through IX [31] (Additional file 6: Figure S1); 2) common NCBI CDS annotations of “serine/threonine protein kinase CDS”; or 3) identification of an STK in previous literature [32]. STKs are mostly associated with eukaryotic signal transduction pathways. In prokaryotes, two-component regulation controls most phosphorylation pathways with a receptor histidine kinase paired with various response regulators phosphorylated on aspartic acid residues. These kinases often control the expression of certain genes [33]. However, Hanks-type STKs have been found in an array of prokaryotic organisms where their genomic abundance is often correlated with genome size, physiological and ecophysiological complexity, and ability to tolerate complex environments [31,32,34]. These STKs are implicated in the regulation of various aspects of bacterial physiology through post-translational modification of proteins, which may themselves be components of phosphorelay and transcriptional regulatory pathways [34–37]. Serine/threonine protein kinases were first associated with the pknA gene of *Nostoc* sp. PCC7120, which is involved in growth and differentiation [32], and in other bacteria their activity regulates processes such as cell growth, segregation, virulence, metabolism, stress adaptation, and cell wall/envelope biogenesis [34]. Ser/Thr kinases in cyanobacteria are usually associated with three different processes: developmental regulation, stress response, and pathogenicity [32]. Slight changes, not in function but in the strength of substrate recognition to a variety of phosphorylation targets, may contribute to the ability to finely tune networks of signal transduction.

Compared to histidine kinases of two-component systems, which exhibit strong substrate discrimination, STKs have relaxed substrate specificities. This has been linked to a lack of co-evolution between the kinase and its cognate target [38,39]. Accelerated evolution of the substrate-binding domain of these kinases may have resulted in the further expansion of this class of proteins in the Cyanobacteria phylum, contributing to a wide range of adaptability to external stimuli and challenging environments. We hypothesize that the VR-containing CLec domain could be autoinhibitory, and activation of kinase activity would occur upon binding a small molecule or protein ligand. In this case, DGR-mediated diversity could allow rapid recognition of various ligands for activating phosphorylation cascades. Alternatively, the CLec domain could function in ligand recognition (i.e. determining what protein(s) are phosphorylated). In this case CLec variants could have different substrate specificities. Segregation of phosphorylation targets between paralogous kinases has been shown to play a strong selective pressure in their evolution [40]. In turn, DGR-driven hypervariation of binding components in signaling proteins may offer additional selective advantages in cyanobacteria through preventing cross-talk, which is characteristic of this of kinase class.

We also identified orthocaspase-like peptidase domains in VP N-termini, which are also common among their paralogs (Fig. 4). Caspase proteins are proteases involved in the initiation of programmed cell death in metazoans [41]. The peptidase domains that we identified in many VPs and their paralogs were predicted as orthocaspases, which are the prokaryotic homologs of eukaryotic caspase-type proteases [42]. While these protein types are homologous to metazoan caspases, current evidence supports a broader role in cell homeostasis during normal cellular conditions, programs of cellular differentiation, or ageing as well as potential apoptosis [43–45]. Previous studies have found orthocaspases to be enriched in morphologically complex filamentous cyanobacteria of subsections III-V (e.g. *Trichodesmium erythraeum* IMS 101, *Anabaena* spp., and *Nostoc* spp.) as well as various strains of the unicellular toxin-producing species, *Microcystis aeruginosa*. Conversely, orthocaspases are entirely absent from unicellular genera *Synechococcus*, *Prochlorococcus*, *Cyanobium*, and *Cyanothece* and are underrepresented in the genomes of cyanobacteria belonging to subsections I-II. This suggests their utility in enabling the complex signal response and regulatory programs that exist in cyanobacteria capable of cellular differentiation, toxin production, and diazotrophy [46].

In addition to the serine/threonine kinase and orthocaspases-like peptidase domains, we identified less-common features including repeat motifs, Toll-like/TIR receptors, GAF, GUN4-like, and CHAT domains. Repeat motifs may have a role in protein-protein interactions (e.g., TPR, WD40 repeat, ARM repeat, and VWA-CoxE) [47–50], while the other domains have been linked to intracellular signal transduction [51–55] (Fig. 5).

A common feature for nearly all of the N-terminal domains, including the prevalent protein kinase-containing paralogs, is their potential to serve a functional role in signal transduction in response to external stimuli (e.g. light, nutrient deprivation, and general stress response) [9]. A previous study found that genes encoding complex multidomain proteins involved in signal transduction are highly enriched in the filamentous cyanobacterium *Anabaena* sp. PCC 7120 when compared to the genomes of unicellular *Synechocystis* sp. PCC 6803 and *Pseudomonas aeruginosa* [56]. Moreover, regulatory proteins involved in signal transduction could lend to the complex regulation necessary for the physiology of filamentous cyanobacteria. These physiologies include a capacity for cell-differentiation, producing heterocysts during nitrogen deprivation and akinetes under environmental stress, as well as programmed apoptosis [43,57]. The presence of DGRs in cyanobacteria follows this trend in the abundance of specialized signal transduction proteins – being seemingly enriched in filamentous nitrogen-fixing taxa and absent from genomes of unicellular taxa, *Synechococcus* spp. and *Prochlorococcus* spp., though they are present in other unicellular species.

DGR-programmed variation of the ligand-binding domain of receptor-binding proteins in Bordetella bacteriophage has been shown to increase the capacity of these proteins to recognize a vast array of molecules. Moreover, diversification of oligomeric structures appeared to confer an amplification of binding affinity, or avidity [58,59]. Specifically, the existence of 12 DGR-variable target protein trimers in each bacteriophage virion was shown to increase the binding strengths of these proteins to their ligand, pertactin, by relaxing the requirement for optimal binding between the ligand and any single monomer. This multivalent binding was also shown to lead to more distinction in binding events, contributing to enhanced selectivity [59]. These two properties of avidity through multivalency are hypothesized to be characteristic of other DGR systems as a means to provide ligand-recognition flexibility to evolve under constrained conditions, while maintaining selectivity. We hypothesize that, in cyanobacteria, DGR-programmed variation might have a role in providing multimeric avidity in terms of ligand binding for signal response. In the case of autoinhibitory variable proteins attached to a kinase, rather than providing flexibility in host-receptor binding, as in Bordetella bacteriophage, increased avidity may hold a kinetic advantage for substrate binding, whereby flexible activation accelerates signal transduction and regulation. More generally, the available genomic evidence is consistent with a phenomenon of targeted diversification acting to tune cyanobacterial regulatory networks.

## Conclusions

The DGR-enabled diversification of proteins involved in host attachment should lead to selective advantages, as this offers an offensive selective advantage to variation by the host cell. By genomic inference, other DGR-containing prokaryotes seem to have adopted DGR function to mechanisms of virulence, and other cell-cell or phage/cell binding interactions. By contrast, our findings suggest a selective use of DGRs for purposes of isolated hyper-diversification of a small pocket in the C-terminal binding domains of multidomain proteins broadly involved in signal transduction within cyanobacteria. This class of DGR-target proteins is thus-far unique to the cyanobacterial phylum. Diversification of the binding site of these proteins, paired with natural selection over iterations of diversity generation and the ability to segregate resulting beneficial mutants via transposition, may contribute to the complexity and adaptability of cellular regulation amongst cyanobacterial taxa. In developing a better grasp on the functional significance of DGR hypervariation, it is clear that the phenomenon adds new layers of complexity in the expansion of bacterial protein networks.

## Methods

### DGR Identification and Annotation

First, we identified all cyanobacterial genomes containing a DGR-RT-like coding sequence by comparing a consensus sequence for previously-identified cyanobacterial DGR-RT sequences against protein databases using pHMMER. All matches were linked to corresponding genome or nucleotide sequences, which were then downloaded from NCBI. A set of potential DGR candidates was first developed using a workflow with Python and Geneious Prime v 2019.2.3 (Biomatters) as previously described [21]. Briefly, RT genes were manually inspected for core NTP-binding site motifs, before searching for near-repeats in a 10-kbp proximal region (i.e., RT +/- 5-kbp). Repeats in this region were then aligned and inspected for: i) random mismatches in one sequence (VR), which predominantly occur in 1^st^ and 2^nd^ codon positions of an ORF, and ii) >80% of mismatches correspond to adenines in the non-coding near-repeat sequence (TR). Next, retroelements were further analyzed using myDGR [60] which is especially effective at identifying putative trans-acting accessory DGR components, and separately, remote VP and VP homolog genes.

The entire DGR dataset contains several RT and VP sequences that are near-identical, but shared by distinct genomes (Additional file 1: Table S1). To generate a representative subset of these redundant DGRs, we used CD-HIT [61] to cluster RT amino acid sequences using the following settings: 0.9 global alignment; 95% identity threshold. For comparison with the global metagenomic DGR dataset developed by Roux et al. [18], we conducted pairwise alignments with RT sequences using blastp [62] and identified similar representatives at ≥ 50% amino acid identity.

Genes, homologous to VPs within the DGR cassette, were inspected by aligning amino acid sequences for the CLec domain of each putative remote VP to the DGR-VP within the same organism using Clustal Omega. Genes with CLec domains having a putative VR with ≥50% nucleotide identity to a DGR-VP were designated as remote VPs, while those with <50% were designated as VP homologs. DGR and remote VP neighborhood regions were defined as regions 10kb upstream and downstream from the DGR cassette, remote VP, or VP homolog.

### Neighborhood Analyses

In order to identify potential transposons, we first examined existing genomic annotations in the neighborhood (i.e. +/- 10 kbp) of each VP and Remote VP for the following features: transposase, integrase, mobile element. Next, we conducted a transposon search using ISFinder [63] using expanded VP loci (60 kbp) that contain one or more annotations associated with mobile elements.

### Phylogenetic Analyses

To construct a phylogenetic tree of cyanobacteria, we used a set of 16 ribosomal proteins often used for phylogenomic analysis (RpL2, 3, 4, 5, 6, 14, 15, 16, 18, 22, and 24, and RpS3, 8, 10, 17, 19 [64]. Each ribosomal protein was identified using HMMER [65] and hidden Markov models from the Pfam [66] database (accessed September 2018). Each individual marker gene was aligned using MUSCLE [67], trimmed using TrimAL [68], manually assessed to remove end gaps and ambiguously aligned regions and concatenated. A maximum likelihood tree was constructed using RAxML v. 8.2.9 [69] with the PROTCATLG model.

To reconstruct RT phylogeny, putative DGR-RT coding sequences were identified, as described above, then translated. Sequences were de-replicated and non-redundant candidates were chosen using CD-Hit [61] with a global alignment threshold of 99% identity. All DGR-RT sequences and a set of Group-II intron RT sequences from Bacteria, Archaea, plastids, and mitochondria were aligned with a hidden markov model of the reverse transcriptase protein family (PF00078) using HMMalign [65]. A phylogenetic tree of DGR-RTs was constructed using FastTree2 [70] with the WAG substitution matrix, and the CAT approximation to optimize branch lengths. The cyanobacterial DGR-RT representatives were extracted from the complete alignment, realigned using Clustal Omega [71] and used to construct an unrooted phylogenetic tree.

### Protein Function Analysis

VP domain architecture was annotated using InterProScan, pHMMER, and HMMScan tools. CD-HIT analysis was performed on CLec domains for all VPs using the following settings: 0.3 global alignment; 30% identity threshold. Amino acid sequences for the CLec domain of all VPs were aligned using Clustal Omega. The C-terminal sequence of all DGR-VP CLecs was extracted based on the InterProScan feature positions, then further aligned using Clustal Omega and a consensus sequence was picked at 75% sequence similarity (Additional File 6: Figure S1). This consensus sequence was used to further identify homologous domains. Using hmmscan, 1,579 hits were returned using an E-value cutoff of 10^−40^ to generate Table S2 (Additional file 2: Table S2).

## Availability of data and materials

The datasets supporting the conclusions of this article are included within the article and its additional files.

## Acknowledgements

This research was funded by a Challenge Grant from the California NanoSystems Institute 439 (CNSI-UCSB). AVE is supported by a National Science Foundation Graduate Research 440 Fellowship Program under Grant No. 1650114, and by the NSF California LSAMP Bridge to the 441 Doctorate Fellowship under Grant No. HRD-1701365. The work conducted by the U.S. 442 Department of Energy Joint Genome Institute is supported by the Office of Science of the U.S. 443 Department of Energy under contract no. DE-AC02-05CH11231. Computational analyses for 444 this work were supported by an NSF-XSEDE resource allocation DEB170007.

## Supplementary information

**Additional File 1: Table S1. RT/Species Table.**

Summary of all DGR-RTs found in cyanobacterial genomes with taxonomic and known physiological information noted. DGR-internal VPs and Remote VPs identified as having VR/TR homology are also noted. See table S6 for information regarding taxonomic affiliations.

**Additional File 2: Table S2. Taxonomy of C-Type Lectin-Like HMM Hits.**

The consensus sequence of cyanobacterial C-type lectin-like predicted domains was generated via alignment of DGR-associated C-termini (up to 200 amino acids) and confirmed with pHMM scan. The taxonomy is summarized for those genomes that contain orthologs.

**Additional File 3: Table S3. Clec Variable Protein Paralogs.**

A subset of 21 high quality genomes were chosen to assess the presence of DGR-VP paralogs. This is a summary of VP paralogs identified and their domains.

**Additional File 4: Table S4. DGR-Proximal Transposable Elements.**

All transposable elements found in proximity to DGRs are listed, including transposase family and mechanisms of integration.

**Additional File 5: Table S5. Genes within DGR Neighborhoods.**

Genes found proximal to cyanobacterial DGR cassettes are annotated with predicted function and counts for each annotated function displayed.

**Additional File 6: Figure S1. Hanks-type Kinase Motif Characterization in VPs.**

Alignment of known “Hanks and Hunter-type” (S/T) kinase domains to the kinase domains of all DGR-VPs, Remote VPs, and VP Paralogs from this dataset. Motifs I-XI highlighted in blue. The top eight sequences denoted “STKII” are known Type II S/T kinases from Zhang et al. 2007 [32].

**Additional File 7: Table S6. Updated Cyanobacterial Taxonomy.**

Taxonomic assignment for each cyanobacterial genome was generated using relative evolutionary divergence and average nucleotide identity with the Genome Taxonomy Database Toolkit (GTDB-Tk) [72] based on the Genome Taxonomy Database (GTDB) [73].

## Notes

### Competing Interest Statement

J.F.M. is a cofounder, equity holder and a member of the Board of Directors of Pylum Biosciences, Inc., a biotherapeutics company in South San Francisco, CA, USA.

