## Supplementary figures and images for "Role of Diversity-Generating Retroelements for Regulatory Pathway Tuning in Cyanobacteria"

### Additional File 6: Figure S1

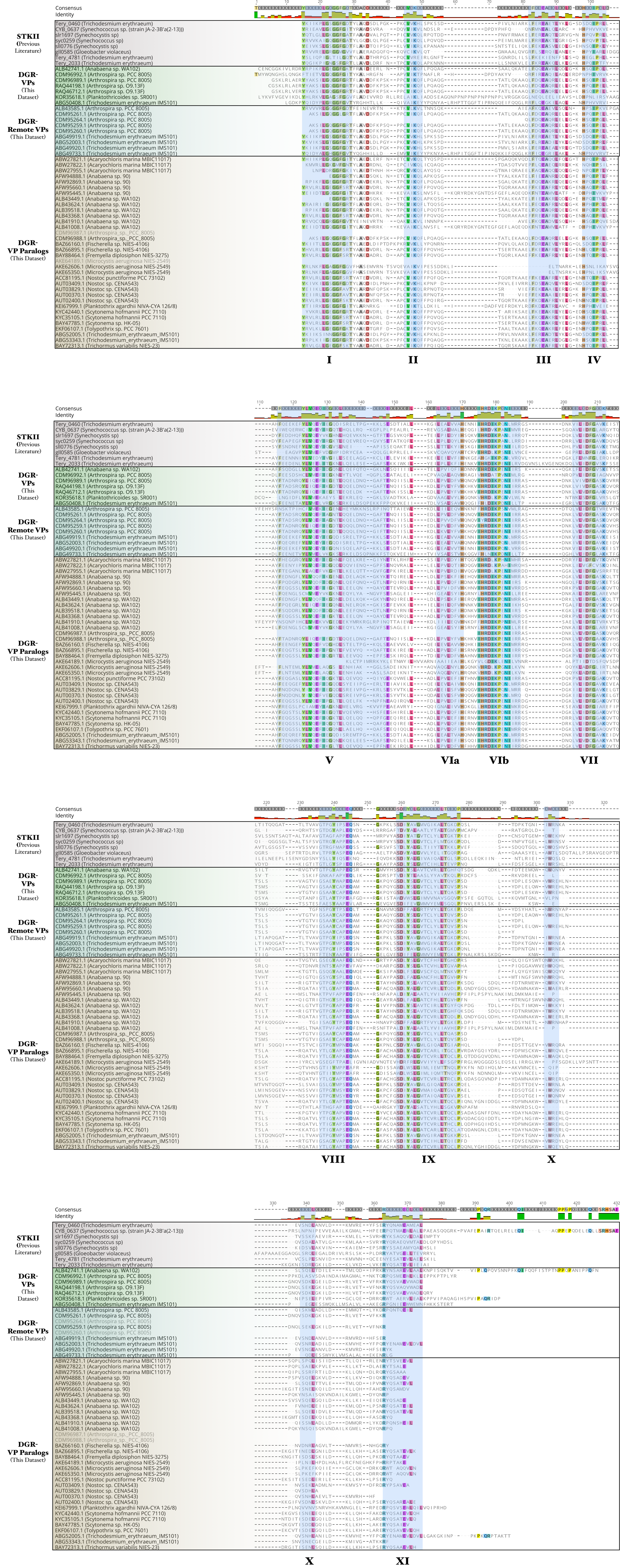
